# Neptune: A Bioinformatics Tool for Rapid Discovery of Genomic Variation in Bacterial Populations

**DOI:** 10.1101/032227

**Authors:** Eric Marinier, Rahat Zaheer, Chrystal Berry, Kelly Weedmark, Michael Domaratzki, Philip Mabon, Natalie Knox, Aleisha Reimer, Morag Graham, Linda Chui, The Canadian Listeria Detection and Surveillance using Next Generation Genomics (LiDS-NG) Consortium, Gary Van Domselaar

## Abstract

The ready availability of vast amounts of genomic sequence data has created the need to rethink comparative genomics algorithms using “big data” approaches. Neptune is an efficient system for rapidly locating differentially abundant genomic content in bacterial populations using an exact *k*-mer matching strategy, while accommodating *k*-mer mismatches. Neptune’s loci discovery process identifies sequences that are sufficiently common to a group of target sequences and sufficiently absent from non-targets using probabilistic models. Neptune uses parallel computing to efficiently identify and extract these loci from draft genome assemblies without requiring multiple sequence alignments or other computationally expensive comparative sequence analyses. Tests on simulated and real data sets showed that Neptune rapidly identifies regions that are both sensitive and specific. We demonstrate that this system can identify trait-specific loci from different bacterial lineages. Neptune is broadly applicable for comparative bacterial analyses, yet will particularly benefit pathogenomic applications, owing to efficient and sensitive discovery of differentially abundant genomic loci.

## Introduction

Capacity to cheaply and quickly generate high volumes of sequence reads has made possible the ability to study the genomes of entire populations of organisms, especially those organisms with relatively small-genomes such as bacteria. Computational biologists have historically used a wide range of bioinformatics software tools to compare small numbers of bacterial genomes and to perform basic characterizations at the nucleotide, gene, and genome scale. However, there now exists a need for bioinformatics software to perform efficient comparative analysis and characterization of entire populations of bacterial genomes. Some tools have emerged. Most of these tools focus on the identification of single nucleotide variants (SNVs) using reference mapping approaches [6, 19], or distance estimations based on small exact substrings (*k*-mers) [10, 15, 22] since these approaches scale well using simple parallelization strategies. Microbial genome-wide association studies (GWAS) that analyze bacterial genome populations to correlate genomic features with phenotypic traits are now also possible thanks to recent methodological developments that address the problems inherent in bacterial genomes that confound conventional GWAS approaches, such as long range linkage disequilibrium and clonal population structure [8]. Some software tools for bacterial GWAS have been developed that associate SNVs or *k*-mers with biological traits [13]. However, for bacterial GWAS, it is important to identify all modes of bacterial genomic variation including larger scale genomic gains and losses, particularly for the majority of bacteria that engage in horizontal gene transfer to acquire novel biologic traits. Scalable software that can rapidly extract the large scale genomic loci that differentiate one population from another while tolerating allelic variation within those loci, is valuable to accomplish bacterial GWAS and has utility for many other applications such as developing targeted molecular diagnostics.

To address this challenge, we looked to the field of genomic signature discovery, where a signature is defined as a genomic locus that is sufficiently represented in a target, or “inclusion” group, and sufficiently absent from a background, or “exclusion” group. An effective signature discovery algorithm is both sensitive and specific, while quick to compute. However, in practice, it remains difficult to develop algorithms that possess all three of these attributes. Early algorithmic approaches for signature discovery were developed with the specific aim of generating pathogen detection diagnostic assays [23]. In general, these approaches involve exhaustively comparing all sequences using alignment-based methods, such as BLAST [1], to locate signature regions in an inclusion group that are absent in the exclusion group. However, these approaches do not scale efficiently and are focused on generating molecular diagnostic primers of a fixed length.

Other more sophisticated approaches attempt to address efficiency by using computationally optimized string processing approaches that encode fixed size substrings from the genome in rapidly searchable data structures, then analyzing these data structures for unique substrings [18]. These approaches are very fast and scale well, but cannot handle variability in the target sequence and are artificially limited to fixed length signatures. Variability in the target can be achieved by grouping similar sequences using multiple sequence alignments [23] or other clustering operations [2]. However, these common clustering techniques come at a high computational cost and do not scale well. Some algorithms incorporate a data reduction step prior to clustering to reduce the amount of unnecessary computation. For example, Insignia [18], TOFI [21], and TOPSI [20] use efficient suffix trees to precompute exact matches within inclusion targets and an exclusion background. However, depending on the size of the background database, this may remain a computationally expensive operation. One interesting novel implementation is CaSSiS [2], which approaches the problem of signature discovery more thoroughly than other signature discovery pipelines. The software produces signatures simultaneously for all locations in a hierarchically clustered data set, such as a phylogenetic tree, thereby producing candidate signatures for all possible subgroups. However, this process requires the input data to be provided in a hierarchically clustered format, such as computationally expensive phylogenies. In addition to the efficiency-versus-sensitivity trade off, most of the programs that have been developed thus far for signature discovery have additional shortcomings that make them unsuitable for identifying common variation between populations of genomes. For example, they may restrict the analysis to a single inclusion genome [21], they don’t permit user-supplied genomes for target identification [18], or they do not provide the software to the end user [23].

We designed Neptune as a system for discovering discriminatory bacterial sequence signatures and conducting comparative analyses of arbitrary groups of genome sequences that leverages existing strategies for signature detection, but in a novel way that is both efficient and accurate. Neptune identifies genomic loci uniquely shared among a user-specified interest group but lacking from a background group. Independent of pre-computation, restriction on targets, and slow clustering approaches, Neptune applies reference-based, parallelized exact-matching *k*-mer strategy for speed, while making allowances for inexact matches to enhance sensitivity. Neptune’s signature discovery is guided with probabilistic models that make decisions with a measure of statistical confidence. Neptune is open-source software freely available at github.com/phac-nml/neptune and is broadly applicable for rapid comparative assessments of bacterial populations.

## Results

### Neptune Design Principles

We define a genomic signature as a string of characters (nucleotides) sufficiently unique to a user-specified set of targets (the inclusion group) that discriminates it from a user-defined background group (the exclusion group). We define a “reference” sequence as any inclusion target from which to extract signatures. Targets will typically comprise draft and closed genome assemblies. Signature discovery aims to locate unique and conserved regions within the inclusion group, but absent or minimally present in the exclusion group.

### Validation

We applied Neptune to identify differentially abundant genomic loci (genomic signatures) for distinct bacterial datasets from broad phyla. In order to validate methodology and highlight mathematical considerations, we first applied Neptune to a simulated *Bacillus anthracis* data set. To demonstrate behaviour in populations with genomic variation dominated by gene gain and loss, we applied Neptune to identify signatures within a clinically-relevant *Listeria monocytogenes* data set. Lastly, we demonstrated Neptune’s capacity to locate genome signatures in a more structurally and compositionally diverse *Escherichia coli* data set.

### Simulated Dataset

In order to show that Neptune identifies signatures as expected, the software was run with an artificially created data set. We created an initial inclusion genome by interspersing non-overlapping, virulence- and pathogen-associated genes from *Vibrio cholerae* M66-2 (NC_012578.1) throughout a *Bacillus anthracis* genome (NC007530) (Table 1). We selected 6 signature regions, varying from 4 kb to 50 kb in size, and spaced these signatures evenly throughout the *B. anthracis* genome with each signature represented only once. The initial exclusion genome consisted of the wildtype *B. anthracis* genome lacking modification. Lastly, we broadened both the inclusion and exclusion groups to 20 genomes each, by generating copies of the corresponding original inclusion or exclusion genome and incorporating a 1% random nucleotide mutation rate, with all possible mutations being equally probable.

**Table 1:** Genomic islands naturally found within *Vibrio cholerae* (NC_012578.1) chromosome I. These islands were used as *in silico* signatures and artificially inserted within a *Bacillus anthracis* genome. These islands were identified with IslandViewer 3 [7].

| ID | Length (bp) | Summary |
| --- | --- | --- |
| 1 | 23338 | O-antigen transport |
| 2 | 50038 | toxin pilus |
| 3 | 12259 | phage replication |
| 4 | 9652 | phage integrase |
| 5 | 4282 | N-acetylneuraminate lyase |
| 6 | 10155 | neuraminidase |

Neptune was used to identify the inserted pathogenic and virulence regions in our simulated *B. anthracis* data set. We specified a *k*-mer size of 27, derived from Equation 2, and used Neptune’s default single nucleotide variant (SNV) rate of 1%. Neptune produced signatures from all 20 inclusion targets (supplementary material) and these signatures were consolidated into a single file. We aligned these signatures to the initial inclusion genome and used GView Server [17] to visualize the identified signatures from all references. Neptune identified 7 consolidated signatures, corresponding to the 6 expected *V. cholerae* regions, with the largest signature region (50 kbp) misreported as two adjacent signatures (10,136 bp and 39,763 bp) with a gap of 143 bp between them. However, by Equation 9, we expect to see erroneous signature breaks with a frequency inversely proportional to our confidence level (95%) when extending signatures over *k*-mer gaps. Indeed, upon investigation, the break location contained six mutations almost evenly spaced within the 143 bp region. Importantly, we observed that all Neptune-identified signatures corresponded to the artificially inserted *V. cholerae* regions and were consistently detected for all references. Neptune reported all of the *in silico* signatures and reported no false positives. Hence, we conclude that Neptune is able to locate all *in silico* signatures; although some regions identified are reported as two adjacent signatures.

#### Listeria monocytogenes

Neptune was used to locate signature regions within two distinct serotypes of *Listeria monocytogenes. L. monocytogenes* is an opportunistic environmental pathogen that causes listeriosis, a serious and life-threatening bacterial disease in humans and animals [16]. *L. monocytogenes* is comprised of a group of genetically heterogeneous strains consisting of clonal isolates with very low recombination rates. However, recent *L. monocytogenes* evolution has been characterized by gene deletion events resulting from horizontally acquired bacteriophage and genomic islands. Hence, we anticipated finding signatures corresponding to these events.

Listeria isolates were serotyped using standard laboratory serotyping procedures [11]. Serotypes 1/2a and 4b were selected for evaluation as they represent distinct evolutionary lineages and are clinically relevant [16]. Of the 13 *L. monocytogenes* serotypes, serotype 1/2b and 4b (lineage I) and serotype 1/2a (lineage II) are most commonly associated with human illness globally [16]. *L. monocytogenes* lineage I is characterized by low diversity and low recombination rates and strains from this lineage are overrepresented amongst human isolates, as compared to lineage II strains, which exhibit increased levels of genomic diversity, owing to recombination and horizontal gene transfer and have an over representation among food, food-related and natural environments [16]. In total, 112 serotype 1/2a and 39 serotype 4b targets were available to be used as inclusion and exclusion groups. These were independently assessed to identify 1/2a signatures as well as the reciprocal 4b signatures, by reversing the inclusion and exclusion groupings. These groups were evenly and randomly subdivided into a test set and a validation set.

Neptune was executed on the *L. monocytogenes* test data in order to produce both 1/2a and 4b signatures for validation. Neptune produced 105 1/2a signatures and 75 4b signatures from their respective inclusion targets. We further evaluated the top-scoring (≥ 0.95) 1/2a and 4b signatures. The top-scoring signatures identified for *L. monocytogenes* serotype 1/2a are listed in Table 2. These signatures included phosphotransferase systems, proteins involved in regulating virulence genes in response to environmental cues, and a surface-exposed internalin protein gene, many of which are known to be critical factors for human pathogenesis [3]. Furthermore, a lineage Il-specific heat-shock system [24], constituting an operon with 3 genes, was present among high scoring signatures. Likewise, the top-scoring signatures identified for *L. monocytogenes* serotype 4b (Table 3) included proteins related to the cell wall, such as teichoic acid biosynthesis and a cell wall anchor protein, and a variety of other signatures encoding broad functional diversity.

**Table 2:** A summary of top-scoring ≥ 0.95 *L. monocytogenes* serotype 1/2a signatures generated by Neptune relative to a serotype 4b background. These signatures were mapped against *L. monocytogenes* 1/2a EGD-e (NC_003210) and 08-5578 (NC_013766) to infer annotations.

| Rank | Score | Length<br>(bp) | Locus Information | *L. monocytogenes*<br>serotype 1/2a str.<br>EGD-e coordinates |
| --- | --- | --- | --- | --- |
| 1 | 0.99 | 4830 | peptidoglycan-bound protein colossin A | 2653185 - 2658013 |
| 2 | 0.99 | 5336 | PTS system, L-ascorbate (L-Asc) family | 2042111 - 2047447 |
| 3 | 0.99 | 4059 | *bvrABC* locus, beta-glucoside-specific sensory system | 2872894 - 2876952 |
| 4 | 0.99 | 5454 | PTS system, glucose-glucoside (Glc) family | 764364 - 769817 |
| 5 | 0.98 | 1938 | hypothetical | 776415 - 778355 |
| 6 | 0.98 | 4514 | two-component response regulator<br>and ABC transport systems | 1086579 - 1091092 |
| 7 | 0.98 | 2839 | internalin | 169228 - 172066 |
| 8 | 0.98 | 1673 | glycosyl-transferase | 532558 - 534230 |
| 9 | 0.97 | 967 | hypothetical | 2717382 - 2718348 |
| 10 | 0.96 | 169 | hypothetical, partial | 270157 - 270325 |
| 11 | 0.96 | 2591 | lineage II specific heat-shock system | 441513 - 444103 |
| 12 | 0.95 | 548 | hypothetical | 804275 - 804822 |

**Table 3:** A summary of top-scoring ≥ 0.95 *L. monocytogenes* serotype 4b signatures generated by Neptune relative to a serotype 1/2a background. The signatures were mapped to *L. monocytogenes* strain 4b F2365 (NC_2973) to infer annotations.

| Rank | Score | Length (bp) | Locus Information | *L. monocytogenes* serotype 4b str. F2365 coordinates |
| --- | --- | --- | --- | --- |
| 1 | 0.99 | 223 | hypothetical | 478246 - 478468 |
| 2 | 0.99 | 3081 | *gltA-gltB* operon | 2787943 - 2791023 |
| 3 | 0.99 | 4004 | N-acetylmuramic acid metabolism | 1685737 - 1689738 |
| 4 | 0.98 | 1709 | cell wall anchor | 2684246 - 2685954 |
| 5 | 0.97 | 1786 | RHS repeat-containing protein (partial) | 471882 - 473667 |
| 6 | 0.97 | 4912 | RHS repeat-containing protein (partial) | 466603 - 471499 |
| 7 | 0.97 | 5917 | multiple, including: hypothetical, cell surface membrane anchor, multidrug efflux transporter like | 428382 - 434298 |
| 8 | 0.97 | 1785 | pyruvyl-transferase | 117970 - 199754 |
| 9 | 0.95 | 1654 | teichoic acid biosynthesis | 2190231 - 2191883 |
| 10 | 0.95 | 1741 | serine protease | 1924193 - 1925933 |

These test-generated signatures were then compared against the wet-lab verified validation data sets to evaluate their *in silico* sensitivity and specificity. We used BLASTN [1] to independently align the top-scoring signatures against our validation data sets. With a percent identity threshold of 95% and a minimum alignment length of 95% the size of the signature length, 670 out of 672 (99.7%) 1/2a signature alignments against the 1/2a validation targets met our sensitivity criteria. Likewise, 199 out of 200 (99.5%) 4b signature alignments against 4b validation targets met this strictness. Similarly, when relaxing the percent identity threshold to 50% and the minimum alignment length to 50% the size of the signature length, we found no 1/2a hits against 4b validation targets and no 4b hits against 1/2a validation targets, indicating that the signatures were specific to the inclusion group. These results suggest that our top-scoring Neptune-identified *L. monocytogenes* serotype 1/2a and 4b signatures have high *in silico* sensitivity and specificity to their respective serotypes against the other serotype background.

#### Escherichia coli

We then applied Neptune to locate signatures corresponding to Shiga-toxin producing *Escherichia coli* (STEC). Specifically, we chose to interrogate *E. coli* genomes that produce the Stx1 toxin. This toxin requires the expression of both the Stx1a and Stx1b subunits to be functional. Therefore, we expected to locate the genes encoding for these subunits using Neptune. As *E. coli* exhibits significantly increased genomic diversity over *L. monocytogenes*, we expect it makes identifying related signatures a more computationally challenging problem.

The inclusion and exclusion data sets were comprised of 6 STEC (Stx1) and 11 non-STEC draft assemblies, respectively (PRJNA57781). Neptune produced 371 signatures corresponding to STEC. The top-scoring signature had nearly 100% *in silico* sensitivity and specificity with respect to the inclusion and exclusion groups. We further investigated the top-scoring (≥ 0.95) consolidated signatures (Table 4) by aligning these signatures against an *E. coli* O157:H7 str. Sakai reference (NC_002695.1, NC_002127.1, NC002128.1) to infer sequence annotations. This alignment included the chromosome and both plasmids, pO157 and pSKA1. The Sakai reference was selected because it contains a copy of the Stx1 toxin and is well characterized.

**Table 4:** A summary of Stxl-containing *E. coli* signatures generated by Neptune relative to a background of non-toxigenic *E. coli*. The signatures were mapped to *E. coli* O157:H7 str. Sakai reference (NC_002695.1, NC_002127.1, NC002128.1) to infer annotations.

| Rank | Score | Length<br>(bp) | Locus Information | *E. coli* O157:H7<br>Sakai coordinates |
| --- | --- | --- | --- | --- |
| 1 | 1.00 | 1375 | Shiga toxin | 2924383 - 2925757 |
| 2 | 0.99 | 5433 | urease gene cluster: *ureA-G* | 1390114 - 1395545 |
| 3 | 0.98 | 3291 | bacteriophage related, integrase and other | 2593022 - 2596313 |
| 4 | 0.98 | 438 | *perC*, transcriptional activator of EaeA/BfpA, partial | 1183201 - 1183639 |
| 5 | 0.98 | 1223 | phage tail length tape measure protein, partial | 2170250 - 2171473 |
| 6 | 0.97 | 7697 | hemolysin gene cluster: *hlyC*, *hlyA*, *hlyB*, *hlyD* | 15716 - 23412 (pO157) |
| 7 | 0.96 | 1260 | colonization factor | 1769157 - 1767898 |
| 8 | 0.96 | 962 | hypothetical | 2200204 - 2201165 |
| 9 | 0.96 | 495 | hypothetical | 2186614 - 2186120 |
| 10 | 0.96 | 796 | phage origin, serine/threonine protein phosphatase | 3488405 - 3489201 |
| 11 | 0.96 | 1364 | hypothetical, colicin-like & small toxic polypeptide | 1397029 - 1398393 |
| 12 | 0.96 | 987 | hypothetical, putative membrane protein | 3486570 - 3487557 |
| 13 | 0.95 | 916 | putative serine acetyltransferase of prophage | 2605160 - 2606076 |
| 14 | 0.95 | 300 | hypothetical, potential T3SS effector | 2209466 - 2209765 |
| 15 | 0.95 | 1136 | T3SS effector protein NleH | 1804974 - 1806122 |

As expected, Neptune identified the Stx1-encoding region as the highest scoring signature (Table 4). Other salient Neptune-identified signatures included several virulence regions such as the urease gene cluster, various phage-related genes, intimin transcription regulator *(perC*) sequences, plasmid-encoded hemolysin gene cluster (Figure 1) and type 3 secretion system (T3SS)-related regions (Table 4).

**Figure 1:**
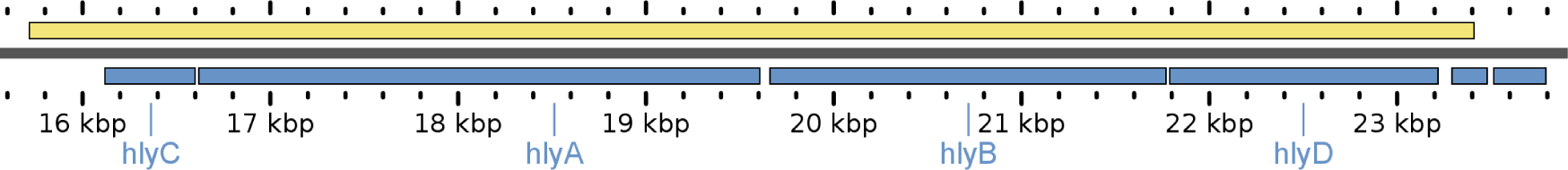
Neptune signature (top) corresponding to a plasmid-encoded *E. coli* hemolysin gene cluster (bottom). GView [17] was used to visualize the 7,697 bp signature above the *E. coli* O157:H7 str. Sakai pO157 plasmid encoding the hemolysin gene cluster.

In the plasmid alignments, the hemolysin-predicted signature was the only top-scoring signature (6th rank; 0.97 score) located on the pO157 plasmid. Furthermore, using BLASTN [1], we found that many of the Neptune top-scoring signatures aligned to characterized *E. coli* O157:H7 O-Islands (a set of mobile genetic islands characterized to carry virulence factors). This included signatures 1-3, 5, 7-15; notably Shiga toxin I (as predicted), a urease gene cluster, and several phage elements. We conclude that Neptune is effective at locating known pathogenic regions and horizontally acquired regions within STEC with high *in silico* sensitivity and high specificity.

## Discussion

### Parameters

While many of Neptune’s parameters are automatically calculated, there are a few parameters that deserve special mention. The minimum number of inclusion hits and maximum gap size are sensitive to the SNV rate and the size of *k*. When estimating these parameters, a slightly higher than expected SNV rate is recommended. This conservative approach will avoid false negatives at the expense of false positives. However, many of these false positives will be removed during the filtering stage at the expense of increased computational time.

### Computation Time

Neptune is parallelizable and performs well on high-performance computing clusters. In order to show the scalability of Neptune, we created a simulated data set by generating one hundred copies of a *L. monocytogenes* serotype 1/2a isolate, one hundred copies of a serotype 4b isolate, and incorporating a 1% random nucleotide mutation rate in each generated copy, with all possible mutations being equally probable. We ran Neptune on a homogeneous computing cluster where there were always more resources available than required by the software. This demonstrates the scalability of Neptune when computing resources are not a limitation (additional information appears in supplementary material). We ran Neptune on 50, 100, 150, and 200 total genomes, with even numbers of inclusion and exclusion genomes, and observed a linear relationship between running time and number of genomes (supplementary material). We observed a relationship suggesting each additional genome added as input would require an additional 10.2s to complete and, more generally, a 53% increase in running time for each additional fold increase in input size.

Neptune may also be run as a parallel process within a single-machine environment. When performing a similar scalability experiment on a smaller real data set comprised of 112 *L. monocytogenes* serotype 1/2a isolates and 38 serotype 4b isolates, run on a single compute node with 48 cores and 80 GB of memory, we observed a linear relationship between the number of genomes and completion time (supplementary material). We varied the size of the input data such that runs maintained an approximate proportion of 70-75% inclusion genomes and 25-30% exclusion genomes. We observed a relationship suggesting each additional genome added as input would require an additional 7.9s to complete and, more generally, a 36% increase in running time for each additional fold increase in input size. The observed difference between this experiment and the previous can be partially attributed to the proportionally smaller exclusion group, from which mutations create more work for the algorithm than from within the inclusion group.

We do not directly compare Neptune against other software, because none of these software are capable of identifying differentially abundant genomic loci with sequence variation, while permitting *ad hoc* input and a variable signature locus size.

### Limitations

Neptune’s signature extraction step avoids false negatives at the expense of false positives. The software attempts to locate signatures that may not contain an abundance of exact matches. This approach produces some false positives. However, false positives are removed during signature filtering and requires increased computational time. As signatures are extracted from a reference, repeated regions do not confound signature discovery. However, if a repeated region is a true signature, then Neptune will report each region as a separate signature. In this circumstance, user curation may be required.

Neptune cannot locate isolated SNVs and other small mutations. Any region with a high degree of similarity to the exclusion group will either not produce candidate signatures or be removed during filtering. Neptune is designed to locate general-purpose signatures of arbitrary size (above a minimum of the *k*-mer size) and does not consider application-specific physical and chemical properties of signatures. While Neptune is capable of producing signatures as small as the *k*-mer size, we observed that very short signatures (< 100 bases) may not contain any seed matches with filtering targets when performing alignments during the filtering process, thereby preventing the signature from being evaluated correctly. We recommend either using smaller seed sizes during pairwise alignments, at the expense of significantly increased computation time, or discretion when evaluating very short signatures.

Finally, Neptune makes assumptions about the probabilistic independence of bases and SNV events; while these events do not occur independently in nature, they allow for significant mathematical simplification. Nonetheless, Neptune is capable of producing highly sensitive and specific signatures using these assumptions.

### Revealing Biology

This study demonstrates that Neptune is a very useful tool for the rapid characterization and classification of pathogenic bacteria of public health significance, as it can efficiently discover differential genomic signatures. Although both *L. monocytogenes* 4b and 1/2a serotypes, belonging to lineages I and II respectively, are associated with human illness, lineage I strains are overrepresented among human cases whereas lineage II isolates are widespread in food-related, natural and farm environments. Among the LiDS-NG project isolates used in our study, 46% of 4b and 17% of 1/2a serotype isolates had a clinical human host origin. Among the signatures for serotype 1/2a, multiple PTS systems and ABC transport systems were found (Table 2 and Supplementary Table 1) which may be correlated to the fact that the presence of a variety of PTS and transport systems provides *L. monocytogenes* serotype 1/2a with a competitive advantage to survive under broad environmental conditions due to its ability to utilize a variety of carbon sources. Among the *L. monocytogenes* 4b serotype signatures found were genes coding for cell-wall anchor proteins, RHS protein known to be associated with mediating intercellular competition and immunity [12], and cell wall polysaccharides and teichoic acid decoration enzymes were found (Table 3 and Supplementary Table 2). In keeping with predilection of lineage I for human clinical disease, such cell-surface components play a role in bacterial-host interactions [4]. The potential involvement of these genes in the virulence and pathogenesis of serotype 4b should be an interesting area of future inquiry.

Interestingly, two very large, but divergent signature sequences corresponding to the 4b (rank 16; score 0.93; length 12685 nt) and 1/2a (rank 14; score 0.94; length 22937) inclusion groups were found by Neptune (Supplementary Table 1 and 2). These serotype-specific signature regions contained non-homologous teichoic acid biosynthesis and transport system genes at equivalent chromosomal locations in the two serotype subgroupings. In addition, another signature (rank 26; score 0.84; length 6348 nt; Supplementary Table 2) spanning 7 genes corresponding to Listeria Pathogenicity Island 3 (LIPI-3, or the listeriolysin S cluster [5]) was only identified in 4b isolates.

In a recent study by Maury *et al.*, (2016) a pattern correlation of gene families with the infection/food ratio of isolates in Listeria pangenome identified virulence associated genes such as LIPI-3 and teichoic acid biosynthesis-related gene clusters in serotype 4b strains to be strongly associated with infectious potential at the population level. Analyzing the same lineage I and II data set analyzed in Maury *et al.*, 2016, Neptune identified signatures that overlapped with our prior, independently generated and distinct genomes for lineage I and II isolates (data not shown).

With the advent of genome-wide association studies and their applications in bacteria to rapidly scan genetic markers as the basis of bacterial phenotypes such as host preference, antibiotic resistance, and virulence across the complete sets of genomes, Neptune offers to be a promising tool to reveal discriminatory genetic markers and associations to particular phenotypic traits. Hence, by generating such a catalogue of differential loci, Neptune is useful in identifying candidate regions for further investigating the association of identified regions with categorical phenotypes, biological traits, or metadata, such as pathogen virulence or persistence in niche environments.

## Conclusion

Neptune allows one to efficiently and rapidly identify genomic loci that are common to one population and distinguishing them from other populations. When applied to pathogens, top-scoring signatures were specific to known regions encoding mobile islands containing pathogenicity-associated coding sequences. While some signatures are reported as smaller, adjacent signatures with intervening gaps, we demonstrated that Neptune can locate signatures in both simulated and biological data sets with high sensitivity and specificity. Neptune provides an array of gene candidates to investigate for their possible role in pathogenesis and functional genomics.

Although Neptune will be useful in broad comparative applications, we anticipate it will be particularly helpful in public health scenarios, where rapid infectious agent screen and characterization is crucial. Neptune may be leveraged to reveal discriminatory signature sequences to uniquely delineate one group of organisms, such as isolates associated with a disease cluster or event, from unrelated sporadic or environmental microbes. Neptune’s computations approach is well suited to comprehensive, *ad hoc* comparisons. We conclude that Neptune is a powerful and flexible tool for locating signature regions with minimal prior knowledge for wide-ranging applications of bacterial characterization.

## Availability

*L. monocytogenes* and *E. coli* data used in the manuscript is stored under NCBI BioProject PRJNA301341. Neptune is developed in Python and the software requires a standard 64-bit Linux environment. The software is available at: *http://github.com/phac-nml/neptune*

## Acknowledgements

The authors would like to thank Franklin Bristow and Eric Enns at The Public Health Agency of Canada for their feedback on various aspects of the software design and implementation. The authors would also like to thank L. Chui, D. Haldane, S. Bekal, and J. Wylie for *E. coli* data.

## Author Contributions

E.M., R.Z., K.W., M.G., G.V.D., and C.B wrote the manuscript. E.M., G.V.D., and M.G. designed the software and E.M. developed the software. E.M. and M.D. designed the mathematical models. E.M., K.W., M.G., and G.V.D. contributed to all experiment design. C.B. and A.R. designed the *L. monocytogenes* experiments. L.C. provided the *L. monocytogenes* data on behalf of the LiDS Consortium. C.B. provided the *E. coli* data. P.M. and N.K. contributed early work on the problem of signature discovery. E.M. performed all experiments. R.Z., E.M., K.W., C.B., G.V.D., and M.G. interpreted the results of the experiments.

## LiDS Consortium

### People

Linda Chui^1^, Laura Patterson-Fortin^1^, Jian Zhang^2^, Franco Pagotto^3^, Jeff Farber^4^, Jim Mahony^5^, Karine Seyer^6^, Sadjia Bekal^7^, Cecile Tremblay^7^, Judy Isaac-Renton^8^, Natalie Prystajecky^8^, Jessica Chen^9^, Morag Graham^10^, Gary Van Domselaar^10^, Natalie Knox^10^, Chrystal Berry^10^, Peter Slade^11^

### Affiliations

^1^University of Alberta, Department of Laboratory Medicine and Pathology, ^2^Alberta Innovates-Technology Futures, ^3^Listeriosi Reference Service, Health Canada, ^4^Health Canada, ^5^McMaster University, Department of Pathology and Molecular Medicine, ^6^Canadian Food Inspection Agency - St. Hyacinthe Laboratory, ^7^Laboratoire de sante publique du Quebec, ^8^BC Public Health & Microbiology Reference Laboratory, ^9^ Univerisity of British Columbia, Department of Food Science, Food, Nutrition and Health, ^10^Public Health Agency Canada-National Microbiology Laboratory,^11^ Maple Leaf Foods

## Methods

Neptune uses the distinct *k*-mers found in each inclusion and exclusion target to identify sequences that are conserved within the inclusion group and absent from the exclusion group. Neptune evaluates all sequence, coding and non-coding, and may therefore produce signatures that correspond to intergenic regions or contain entire operons. The *k*-mer generation step produces distinct *k*-mers from all targets and aggregates this information, reporting the number of inclusion and exclusion targets that contain each *k*-mer. The signature extraction step identifies candidate signatures from one or more references, which are assumed to be drawn from inclusion targets. Candidate signatures are filtered by performing an analysis of signature specificity using pairwise sequence alignments. The remaining signatures are ranked by their Neptune-defined sensitivity and specificity scores, representing a measure of signature confidence.

We provide descriptions of the different stages of signature discovery below and an overview of the signature discovery process is found in Figure 2. The majority of parameters within Neptune are automatically calculated for every reference. However, the user may specify any of these parameters. A full description of the mathematics used in the software is provided in the supplementary materials. In our probabilistic model, we assume that the probability of observing any single nucleotide base in a sequence is equal to and independent of all other positions and the probability of all single nucleotide variant (SNV) events (e.g. mutations, sequencing errors) occurring is equal to and independent of all other SNV events.

**Figure 2:**
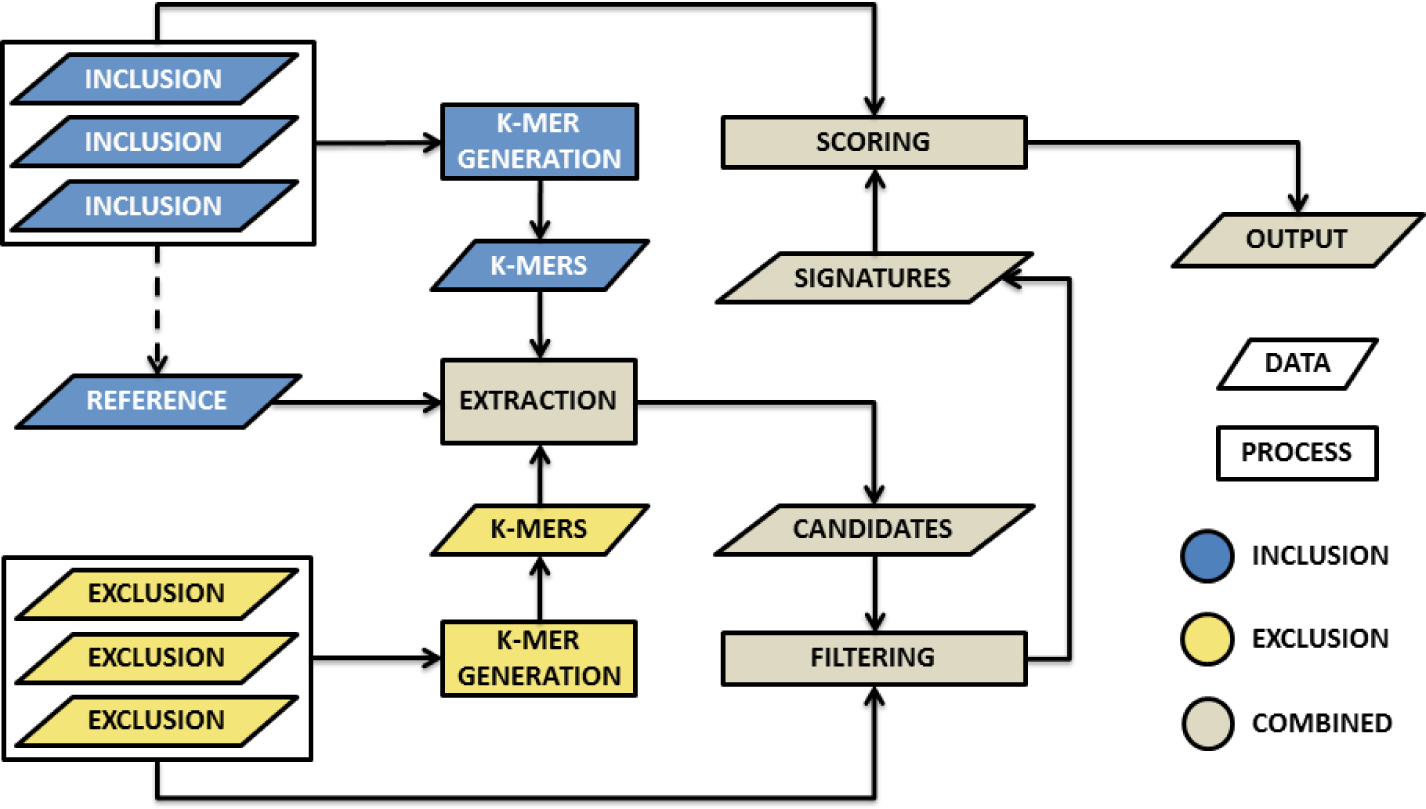
An overview of Neptune’s signature discovery process for a single target. The first step involves generating *k*-mers from all inclusion and exclusion targets. These *k*-mers are aggregated and provided as input to signature extraction. Signature extraction produces candidate signatures, which are filtered and then sorted by their sensitivity and specificity scores.

### *k*-mer Generation

Neptune produces the distinct set of *k*-mers for every inclusion and exclusion target and aggregates these *k*-mers together before further processing. The software is concerned only with the existence of a *k*-mer within each target and not with the number of times a *k*-mer is repeated within a target. Neptune converts all *k*-mers to the lexicographically smaller of either the forward *k*-mer or its reverse complement. This avoids maintaining both the forward and reverse complement sequence [14]. The number of possible *k*-mers is bound by the total length of all targets. The *k*-mers of each target are determined independently and, when possible, in parallel. In order to facilitate parallelizable *k*-mer aggregation, the *k*-mers for each target may be organized into several output files. The *k*-mers in each file are unique to one target (e.g., isolate genome or sequence) and all share the same initial sequence index. This degree of organization may be specified by the user.

The *k*-mer length is automatically calculated unless provided by the user. A summary of recommended *k*-mer sizes for various genomes can be found in Supplementary Table 1. We suggest a size of *k* such that we do not expect to see two arbitrary *k*-mers within the same target match exactly. This recommendation is motivated by wanting to generate distinct *k*-mer information, thereby having matching *k*-mers most often be a consequence of nucleotide homology. Let λ be the most extreme GC-content of all targets and ω be the size of the largest target in bases. The probability of any two arbitrary *k*-mers, *k_X_* and *k_Y_*, matching exactly, *P*(*k*_*X*_ = *k_Y_*)_*A*_, where *x* ≠ *y*, is defined as follows:

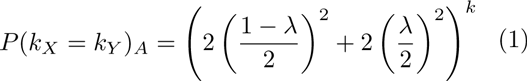

We use the probability of arbitrary *k*-mers matching, *P*(*k*_*X*_ = *k*_*Y*_)_*A*_, to approximate the probability of *k*-mers matching within a target, *P*(*k_X_* = *k_Y_*). This is an approximation because the probability of *P*(*k*_*X*+1_ = *k*_*Y*+1_) is known to not be independent of *P*(*k_X_* = *k_Y_*). However, this approximation approaches equality as *P*(*k*_*X*_ = *k*_*Y*_)a decreases, which is accomplished by selecting a sufficiently large *k*, such that we do not expect to see any arbitrary *k*-mer matches. We suggest using a large enough *k* such that the expected number of intra-target *k*-mer matches is as follows:

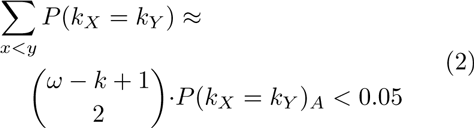

The distinct sets of *k*-mers from all targets are aggregated into a single file, which is used to inform signature extraction. This process may be performed in parallel by aggregating *k*-mers sharing the same initial sequence index and concatenating the aggregated files. Aggregation produces a list of *k*-mers and two values (the number of inclusion and exclusion targets containing the *k*-mer, respectively). This information is used in the signature extraction step to categorize some *k*-mers as inclusion or exclusion *k*-mers.

### Extraction

Signatures are extracted from one or more references, which are drawn from all inclusion targets, unless specified otherwise. However, our probabilistic model assumes all references are included as inclusion targets. In order to identify candidate signatures, Neptune reduces the effective search space of signatures by leveraging the spatial sequencing information inherent within the references. Neptune evaluates all *k*-mers in each reference, which may be classified as inclusion or exclusion *k*-mers. An inclusion *k*-mer is observed in a sufficient number of inclusion targets and not observed in a sufficient number of exclusion targets. The sufficiency requirement is described below. Inclusion and exclusion *k*-mers are used to infer inclusion and exclusion sequence, with signatures containing primarily inclusion sequence. An inclusion *k*-mer may contain both inclusion and exclusion sequence because, while they may contain exclusion sequence, *k*-mers that overlap inclusion and exclusion sequence will often be unique to the inclusion group. An exclusion *k*-mer is, by default, any *k*-mer that has been observed at least once in any exclusion target. However, in some applications it may be desirable to relax this stringency. For example, leniency may be appropriate when the inclusion and exclusion groups are not fully understood. This may be the case when meta data is incomplete or unreliable. An exclusion *k*-mer should, by design, not contain any inclusion sequence. Neptune outputs several “candidate signatures”, which begin with the last base position of the first inclusion *k*-mer, contain an allowable number of *k*-mer gaps and no exclusion *k*-mers, and end with the first base position of the last inclusion *k*-mer (Figure 3). This process is conceptually similar to taking the intersection of inclusion *k*-mers and allowable *k*-mer gaps. Furthermore, it avoids generating a candidate containing exclusion sequence found in inclusion *k*-mers that overlap inclusion and exclusion sequence regions.

**Figure 3:**
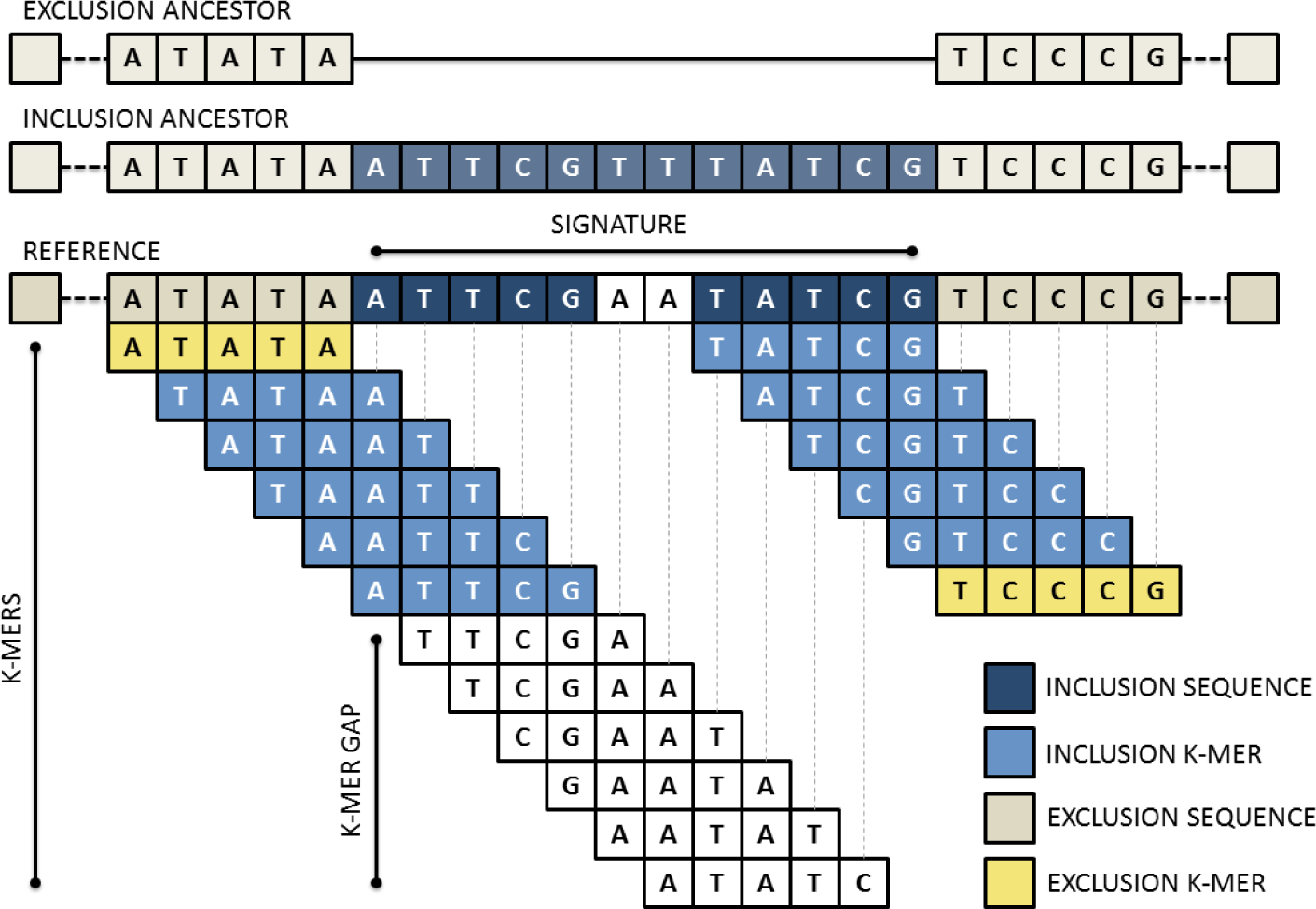
An overview of Neptune’s signature extraction process. The reference is decomposed into its composite *k*-mers. These *k*-mers may be classified as either inclusion or exclusion and are used to infer inclusion and exclusion sequence in the reference. A signature is constructed from inclusion *k*-mers containing sufficiently small *k*-mer gaps and no exclusion *k*-mers.

An inclusion *k*-mer is considered sufficiently represented when it is observed in a number of targets exceeding a minimum threshold. We assume that if there is a signature present in all inclusion targets, then the signature will correspond to homologous sequences in all these targets and these sequences will produce exact matching *k*-mers with some probability. We start with the probability that two of these homologous bases, *X* and *Y*, match is:

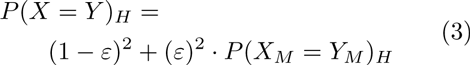

where *ε* is the probability that two homologous bases do not match exactly, and *P*(*X*_*M*_ = *Y*_*M*_)_*H*_ is the probability that two homologous bases both mutate to the same base. The default probability of *ε* is 0.01. We assume that when the homologous bases do not match, the observed base is dependent on the GC-content of the environment. Let λ be the GC-content of the environment. The probability of *P*(*X*_*M*_ = *Y*_*M*_)_*H*_ is defined as follows:

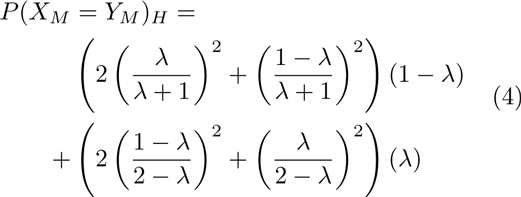

This probability depends significantly on GC-content of the environment. We assume that the probability of each base matching is independent. Therefore, the probability that two homologous *k*-mers, *k*_*X*_ and *k*_*Y*_, match:

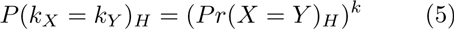

We model the process of homologous *k*-mer matches with a binomial distribution. If we are observing a true signature region in a reference, we expect that corresponding homologous *k*-mers exist in all inclusion targets and infer this homology from aggregated *k*-mer information. An observed reference *k*-mer will exactly match a corresponding homologous *k*-mer in another inclusion target with a probability of *p* = *P*(*k*_*X*_ = *k*_*Y*_)_*H*_ and not match with a probability of *q* = 1 − *p*. The expected number of exact *k*-mer matches with a reference *k*-mer will be *μ* = (*n* − 1) · *p* and the variance will be *σ*^2^ = (*n* − 1) · *p* · *q*, where *n* is the number of inclusion targets. We require *n* − 1 because the reference is an inclusion target and its *k*-mers will exactly match themselves. However, we compensate for this match in our expectation calculation. We assume the probability of each *k*-mer match is independent and that *k*-mer matches are a consequence of homology. When the number of inclusion targets and the probability of homologous *k*-mers exactly matching are together sufficiently large, the binomial distribution is approximately normal. Let *α* be our statistical confidence and Φ^−1^(α) be the probit function. The minimum number of inclusion targets containing a *k*-mer, Λ_in_, required for a reference *k*-mer to be considered an inclusion *k*-mer is defined as follows:

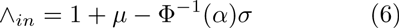

The Λ_*in*_ parameter is automatically calculated unless provided by the user and will inform candidate signature extraction. However, there may be mismatches in the reference, which exclude it from the largest homologous *k*-mer matching group. We accommodate for this possibility by allowing *k*-mer gaps in our extraction process. We model the problem of maximum *k*-mer gap size between exact matching inclusion *k*-mers as recurrence times of success runs in Bernoulli trials. The mean and variance of the distribution of the recurrence times of *k* successes in Bernoulli trials is described in Feller 1960 [9]:

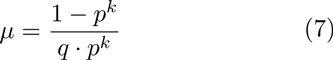

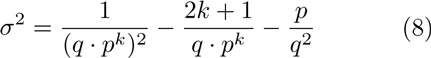

This distribution captures how many bases we expect to observe before we see another homologous *k*-mer match. The probability of a success is defined at the base level as *p* = *P*(*X* = *Y*)_*H*_ and the probability of failure as *q* = (1 − *p*). This distribution may not be normal for a small number of observations. However, we can use Chebyshev’s Inequality to make lower-bound claims about the distribution:

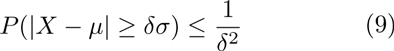

where *δ* is the number of standard deviations, *σ*, from the mean, *μ*. Let *P*(|*X* − *μ*| ≥ *δσ*) be our statistical confidence, *α*. The maximum allowable *k*-mer gap size, V_*gap*_, is calculated as follows:

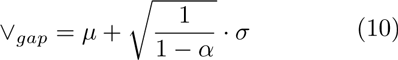

The V_*gap*_ parameter is automatically calculated unless specified. Candidate signatures are terminated when either no additional inclusion *k*-mers are located within the maximum gap size, V_*gap*_, or an exclusion *k*-mer is identified. In both cases, the candidate signature ends with the last inclusion *k*-mer match. The consequence of terminating a signature early is that a large, contiguous signature may be reported as multiple smaller signatures. We require the minimum signature size, by default, to be four times the size of *k*. However, for some applications, such as designing assay targets, it may be desirable to use a smaller or larger minimum signature size. Signatures cannot be shorter than *k* bases. We found that smaller signatures were more sensitive to the seed size used in filtering alignments. There is no maximum signature size. As a consequence of Neptune’s signature extraction process, signatures extracted from the same target may never overlap each other.

### Filtering

The candidate signatures produced will be relatively sensitive, but not necessarily specific, because signature extraction is done using exact *k*-mer matches. The candidate signatures are guaranteed to contain no more exact matches with any exclusion *k*-mer than was specified in advanced by the user. However, there may exist inexact matches within exclusion targets. Neptune uses BLAST [1] to locate signatures that align with any exclusion target and, by default, removes any signature that shares 50% identity with any exclusion target aligning to at least 50% of the signature, anywhere along the signature. This process is done to avoid investigating signatures that are not highly discriminatory. The remaining signatures are considered filtered signatures and are believed to be sensitive and specific, within the context of the relative uniqueness of the input inclusion and exclusion groups, and the parameters supplied for target identification.

### Scoring

Signatures are assigned an overall score corresponding to their highest-scoring BLAST [1] alignments with all inclusion and exclusion targets. This score is the sum of a positive inclusion component and a negative exclusion component, which are analogous to sensitivity and specificity, respectively, with respect to the input data. Let |*A*(*S*, *I*_*i*_)| be the length of the highest-scoring aligned region between a signature, *S*, and an inclusion target, *I*_*i*_. Let |*S*| be the length of signature *S*, *PI*(*S*, *I*_*i*_) the percent identity (identities divided by the alignment length) between the aligned region of *S* and *I*_*i*_, and |*I*| be the number inclusion targets. The negative exclusion component is similarly defined. The signature score, *score*(*S*), is calculated as follows:

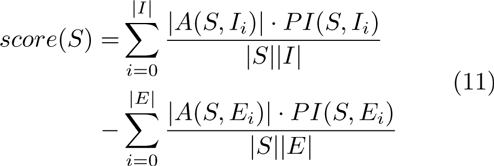

This score is maximized when all inclusion targets contain a region exactly matching the entire signature and there exists no exclusion targets that match the signature. Signatures are sorted based on their scores with highest-ranking signatures appearing first in the output.

### Output

Neptune produces a list of candidate, filtered, and sorted signatures for all references. The candidate signatures are guaranteed to contain, by default, no exact matches with any exclusion *k*-mer. However, there may still remain potential inexact matches within exclusion targets. The filtered signatures contain no signatures with significant sequence similarity to any exclusion target. Sorted signatures are filtered signatures appearing in descending order of their signature scores. A consolidated signature file is additionally provided as part of Neptune’s output. This file contains a consolidated list of the top-scoring signatures produced from all reference targets, such that homologous signatures are reported only once. However, because this file is constructed in a greedy manner, it is possible for signatures within this file to overlap each other. To identify redundancy across the reference targets, we recommend evaluating the signatures identified from each individual reference target in combination with this consolidated file when evaluating signatures.

